# Homeostatic regulation of axonal K_v_1.1 channels accounts for both synaptic and intrinsic modifications in CA3 circuit

**DOI:** 10.1101/2021.04.22.440937

**Authors:** Mickaël Zbili, Sylvain Rama, Maria-José Benitez, Laure Fronzaroli-Molinieres, Andrzej Bialowas, Norah Boumedine-Guignon, Juan José Garrido, Dominique Debanne

## Abstract

Homeostatic plasticity of intrinsic excitability goes hand-in-hand with homeostatic plasticity of synaptic transmission. However, the mechanisms linking the two forms of homeostatic regulation have not been identified so far. Using electrophysiological, imaging and immunohistochemical techniques, we show here that blockade of excitatory synaptic receptors for 2-3 days induces an up-regulation of synaptic strength at CA3-CA3 connexions and intrinsic excitability of CA3 pyramidal neurons. Activity-deprived connexions were found to express a high release probability, an insensitivity to dendrotoxin, and a lack of depolarization-induced presynaptic facilitation, indicating a loss of presynaptic Kv1.1 function. The down-regulation of Kv1.1 channels in activity-deprived neurons was confirmed by their broader action potentials measured in the axon that were insensitive to dendrotoxin. We conclude that regulation of axonal Kv1.1 channel constitutes a unique mechanism linking intrinsic excitability and synaptic strength that accounts for the functional synergy existing between homeostatic regulation of intrinsic excitability and synaptic transmission.

## Introduction

Chronic modulations of activity regime in neuronal circuits induce homeostatic plasticity. This implicates a regulation of both intrinsic excitability (homeostatic plasticity of intrinsic excitability; HP-IE) and synaptic transmission (homeostatic plasticity of synaptic transmission; HP-ST) to maintain networks activity within physiological bounds (*1*). In most cases, these two forms of homeostatic plasticity act synergistically but involve different molecular actors. HP-IE involves the regulation of voltage-gated ions channels (*2–9*) while HP-ST involves the regulation of postsynaptic receptors to neurotransmitters (*10–17*) or the regulation of the readily releasable pool of synaptic vesicles (*18–20*). However, the function of voltage-gated ion channels is not limited to the control of intrinsic excitability (IE). Several studies point the role of axonal voltage-gated channels in shaping presynaptic action potential (AP) waveform and subsequently controlling neurotransmitter release and synaptic transmission (ST) (*21*– *35*). Moreover, some studies describe homeostatic plasticity of AP waveform via voltage-gated channels regulation (*36–38*) while other studies report an absence of this phenomenon (*39*).

K_v_1.1 channel are responsible of the fast activating, slow inactivating D-type current (I_D_) in CA3 neurons. This current has been shown to create a delay in the onset of the first AP and to determine IE in various neuronal types, including CA1 and CA3 pyramidal neurons of the hippocampus (*3, 40*), L5 pyramidal neurons of the cortex (*26, 34*) and L2-3 fast-spiking interneurons of the somatosensory cortex (*41, 42*). Furthermore, K_v_1.1 channels have been shown to control axonal AP width and subsequently presynaptic calcium entry and neurotransmitter release. In fact, pharmacological suppression of K_v_1.1 channels broaden presynaptic AP and increase ST at neocortical and hippocampal glutamatergic synapses and at cerebellar GABAergic synapses (*21, 22, 26, 30, 32, 41, 43, 44*). Moreover, K_v_1.1 channels have been shown to be involved in the phenomenon of depolarization induced Analog Digital Facilitation of synaptic transmission (d-ADF). In fact, at CA3-CA3 and L5-L5 synapses, a somatic subthreshold depolarization of the presynaptic cell leads to inactivation of axonal K_v_1.1 channels, inducing the broadening of the presynaptic AP, an increase in spike-evoked calcium entry and a facilitation of presynaptic glutamate release (*26, 31, 33, 34, 45, 46*). Therefore, Kv1.1 channels control both IE and glutamate release in CA3 pyramidal neurons.

K_v_1.1 channels have been shown to be involved in HP-IE. Chronic activity enhancement by kainate application leads to an increase in I_D_ current and a decrease in IE in Dentate Gyrus granule cells (*47*). Conversely, chronic sensory deprivation leads to K_v_1.1 channels down-regulation and enhancement of IE in the avian cochlear nucleus (*48*). In the CA3 hippocampal network, chronic activity deprivation entails both a homeostatic increase in CA3 pyramidal neurons IE (*3*) and a homeostatic enhancement of ST between pairs of CA3 neurons (*20*). Therefore, the HP-IE and the HP-ST act synergistically to maintain the activity of the CA3 network within physiological bounds. The HP-IE has been shown to involve the downregulation of K_v_1.1 channels (*3*). In this study, we examined whether the increase in ST is also due to K_v_1.1 channel downregulation which would possibly explain the observed synergy between HP-IE and HP-ST.

We show here that chronic AD induced with an antagonist of ionotropic glutamate receptors (kynurenate) in hippocampal organotypic cultures provokes both an increase in CA3 pyramidal cells excitability (HP-IE) and an enhancement of synaptic strength between monosynaptically connected CA3 neurons (HP-ST). Moreover, AD cultures display a decrease in K_v_1.1 channels staining in the axon initial segment but an increase in K_v_1.1 channels staining in the soma of CA3 pyramidal cells. Consistent with this result, bath application of DTX-K, a selective blocker of K_v_1.1 channels, leads to a larger increase in IE in control cultures than in AD cultures. Moreover, focal puffing of DTX-K on axon increases IE in control but not in AD cultures, showing that HP-IE in AD cultures is partly due to the downregulation of axonal K_v_1.1 channels. In addition, we showed that axonal K_v_1.1 downregulation in AD cultures is also responsible for a spike broadening in CA3 neurons, leading to an increase in release probability at CA3-CA3 synapses. Consistent with that, d-ADF, a K_v_1.1-dependent synaptic facilitation, is present in control cultures but not in AD cultures. Altogether, these results show that chronic activity blockade of the CA3 network induces the downregulation of axonal K_v_1.1 channels leading to a homeostatic increase in both IE and ST, thus accounting for the synergistic effect of HP-IE and HP-ST.

## Results

### Increased IE and ST in activity deprived CA3 circuits

To determine the effect of chronic activity deprivation on the CA3 hippocampal network, we blocked excitatory synaptic transmission in hippocampal organotypic cultures (8-12 DIV) with kynurenate (2 mM) for 2-3 days. We found an increase in CA3 pyramidal cell IE in activity-deprived (AD) cultures compared to control cultures (**Figure 1A)**. In fact, the rheobase of CA3 pyramidal neurons was reduced in AD cultures (control: 102.3 ± 7.4 pA, n = 13; AD: 60 ± 7 pA, n = 12; Mann-Whitney test: p < 0.01; **Figure 1B, left**) and the gain of their input/output curve was increased in AD cultures (control: 0.109 ± 0.013 spikes/pA, n = 13; AD: 0.137 ± 0.013 spikes/pA, n = 12; Mann-Whitney test: p < 0.05; **Figure 1B, right**). In order to evaluate the effects of activity deprivation on synaptic transmission, we performed recordings of monosynaptically connected pairs of CA3 pyramidal neurons. The postsynaptic neurons were recorded in voltage-clamp (**Figure 1C**) or in current-clamp (**Figure 1D**) and EPSC/Ps were compared in the two conditions. In AD organotypic cultures, we found an increase in both EPSCs amplitude (control cultures: -25.47 ± 1.98 pA, n = 103; AD cultures: -34.64 ± 3.93 pA, n = 42, Mann-Whitney test: p<0.05; **Figure 1C**) and EPSPs amplitude (control cultures: 1.69 ± 0.23 mV, n = 60; AD cultures: 3.15 ± 0.74 pA, n = 23, Mann-Whitney test: p<0.05; **Figure 1D**) at synapses between CA3 pyramidal neurons. Contrary to previously reported results (*20*), we did not find a significant decrease in the connectivity between CA3 pyramidal neurons in AD cultures (control: 48% of connected pairs among 341 tested pairs; AD: 39% of connected pairs among 165 tested pairs, χ^2^ test: p>0.05; **Supplementary Figure 1**). We can conclude from these results that a chronic activity deprivation by kynurenate treatment induces homeostatic compensations of both intrinsic excitability (HP-IE) of CA3 pyramidal neurons and synaptic transmission (HP-ST) between CA3 pyramidal neurons.

**Figure 1:**
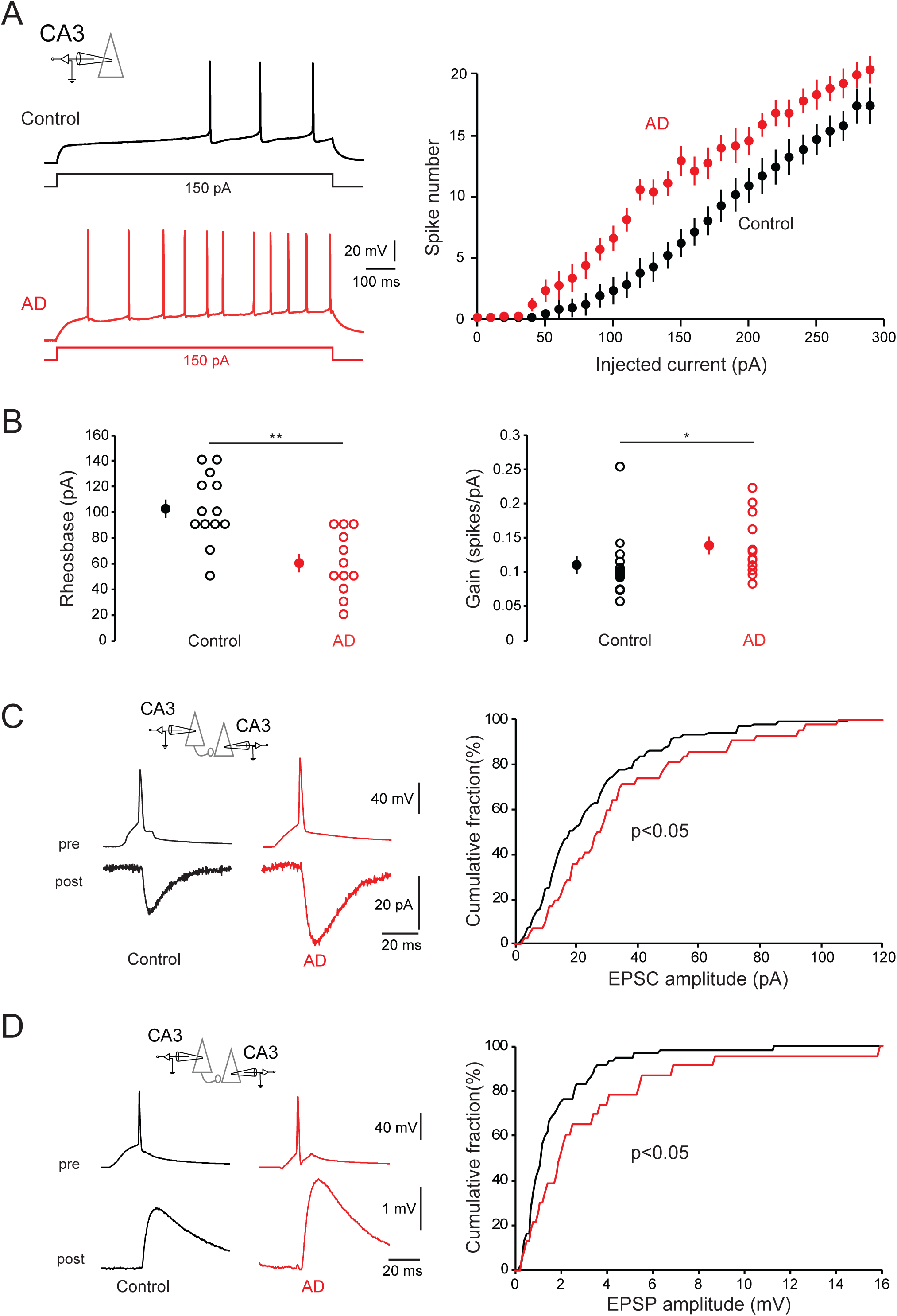
Increase in IE and ST in CA3 circuit following chronic AD. A. Increase in IE in activity deprived (AD) cultures (red) compared to control cultures (black). Left, Example current-clamp traces recorded in CA3 pyramidal neurons in control (black) and in AD (red) cultures. Right, Average data across groups. CA3 pyramidal neurons display a larger excitability in AD cultures (red; n = 13) than in control cultures (black; n = 12). B. Left, Reduced rheobase in AD cultures (red) compared to control cultures (black). Right, Increased input/output curve gain in AD cultures (red) compared to control cultures (black). C-D. Increase in synaptic strength in activity deprived (AD) cultures (red) compared to control cultures (black). C. Left, examples of CA3-CA3 EPSCs in control and AD cultures (average of 30 traces). Right, cumulative histogram of EPSC amplitude in control (black) and AD (red) cultures. Note the rightward shift for AD cultures showing a global increase in EPSC amplitude. D. Left, examples of CA3-CA3 EPSPs in control and AD cultures (average of 30 traces). Right, cumulative histogram of EPSP amplitude in control (black) and AD (red) cultures. Note the rightward shift for AD cultures showing a global increase in EPSP amplitude.

### Reduced axonal K_v_1.1 channels density in AD cultures

It was previously shown that a kynurenate treatment for 2-3 days decreases I_D_ current in CA3 pyramidal neurons (*3*). Therefore, we performed co-immunostaining of K_v_1.1 channels and AnkyrinG in control and AD cultures (**Figure 2)**. In control cultures, we found K_v_1.1 channels staining colocalized with Ankyrin G in the distal part of the axon initial segments (AIS) of CA3 pyramidal neurons (**Figure 2A)**. In AD cultures, we found a 140% increase in the K_v_1.1 channels staining in the somas (Mann-Whitney test: p<0.01; **Figure 2A-B-C**) and a 45% decrease in K_v_1.1 channels staining in the AIS (Mann-Whitney test: p<0.01; **Figure 2A-D-E**), suggesting an accumulation of Kv1.1 channels in the soma probably due to reduced transport of Kv1.1 channels to the axon. Moreover, we found a 40% increase of AIS length in AD cultures (control: 29.99 ± 0.29 μm, n = 100; AD: 42.18 ± 0.40 μm, n = 100; Mann-Whitney: p < 0.0001; **Figure 2F**). These results are similar to the AIS length increase and the K_v_1.1 decrease observed in nucleus magnocellularis (NM) neurons after auditory inputs deprivation (*48*). However, we did not find any modification in the AIS starting position in AD cultures (data not shown). We conclude that chronic activity deprivation leads to a decrease in K_v_1.1 channels density at the AIS and an increase of AIS length in CA3 pyramidal neurons.

**Figure 2:**
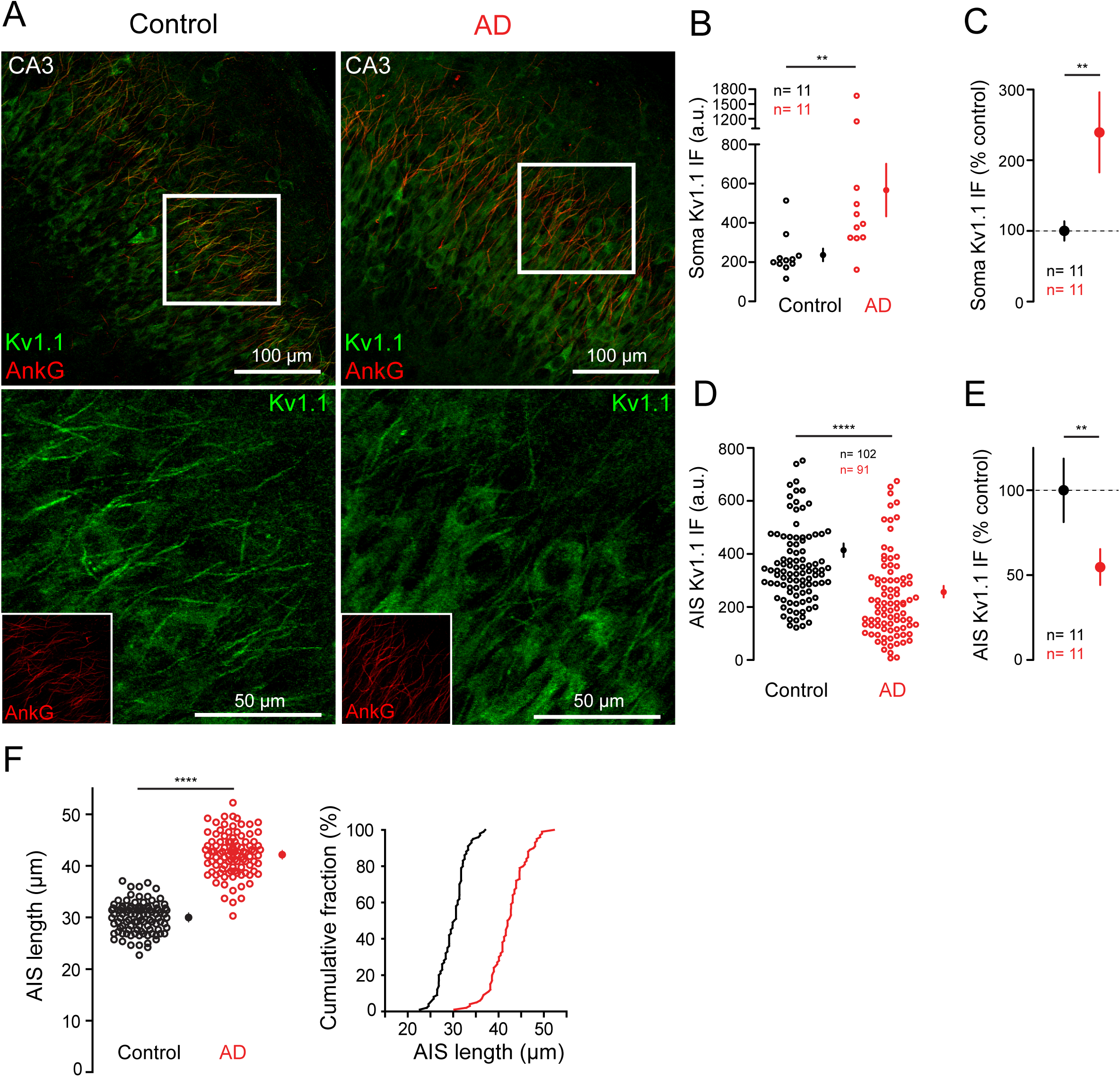
Decrease in axonal K_v_1.1 expression in AD cultures. A. Hippocampus CA3 region of control and AD hippocampal slice cultures stained with K_v_1.1 (green) and Ankyrin G (red) antibodies (upper panels). Lower panels show an amplified image of K_v_1.1 staining in the region indicated in upper panels (box). Ankyrin G staining of this region is included in their respective lower panels. Note the decreased K_v_1.1 expression at AIS accompanied by an increase in neuronal somas in activity deprived cultures. B. K_v_1.1 integrated fluorescence intensity in neuronal somas of each slice (a.u.). C. Mean K_v_1.1 immunofluorescence intensity in neuronal somas per slice. The values are normalized by the mean value in control slices. D. K_v_1.1 integrated fluorescence intensity in axon initial segments (a.u.). E. Mean K_v_1.1 immunofluorescence intensity in AIS per slice. The values are normalized by the mean value in control slices. F. Left, AISs length in control and AD slices. Right, cumulative frequency distribution of AIS lengths in control and AD slices.

### Increased IE in AD cultures is due to reduction of axonal K_v_1.1 channels

In order to observe if HP-IE is due to axonal K_v_1.1 channels density decrease, we performed bath application of DTX-K, a K_v_1.1 channels blocker. K_v_1.1 channels blockade led to a larger increase in IE in CA3 pyramidal neurons from control cultures than from AD cultures (**Figure 3A-B**). In fact, the rheobase reduction following DTX-K application was larger in control cultures than in AD cultures (control: -46.7 ± 9.4 pA, n = 9; AD: -19 ± 5.7 pA, n = 10; Mann-Whitney test: p < 0.05; **Figure 3B, left**). However, DTX-K application led to a similar increase in the input/output curve gain in control cultures and in AD cultures (control: 0.105 ± 0.039 pA, n = 9; AD: 0.072 ± 0.015 pA, n = 10; Mann-Whitney test: p > 0.1; **Figure 3B, right**). Importantly, the intrinsic excitability of CA3 pyramidal neurons was similar in control and AD cultures after DTX-K application (**Supp Fig.2, left**). In fact, the rheobase was similar in control and AD cultures after DTX-K application (control: 54.4 ± 9.3 pA, n = 9; AD: 41.0 ± 6.2 pA, n = 10; Mann-Whitney test: p > 0.1; **Supp Fig.2, middle**). Similarly, the input/output curve gain was similar in control and AD cultures after DTX-K application (control: 0.224 ± 0.042 pA, n = 9; AD: 0.212 ± 0.018 pA, n=10; Mann-Whitney test: p > 0.1; **Supp Fig.2, right**). Therefore, we conclude that the homeostatic increase in IE in AD neurons was due to the downregulation of Kv1.1 channels.

**Figure 3:**
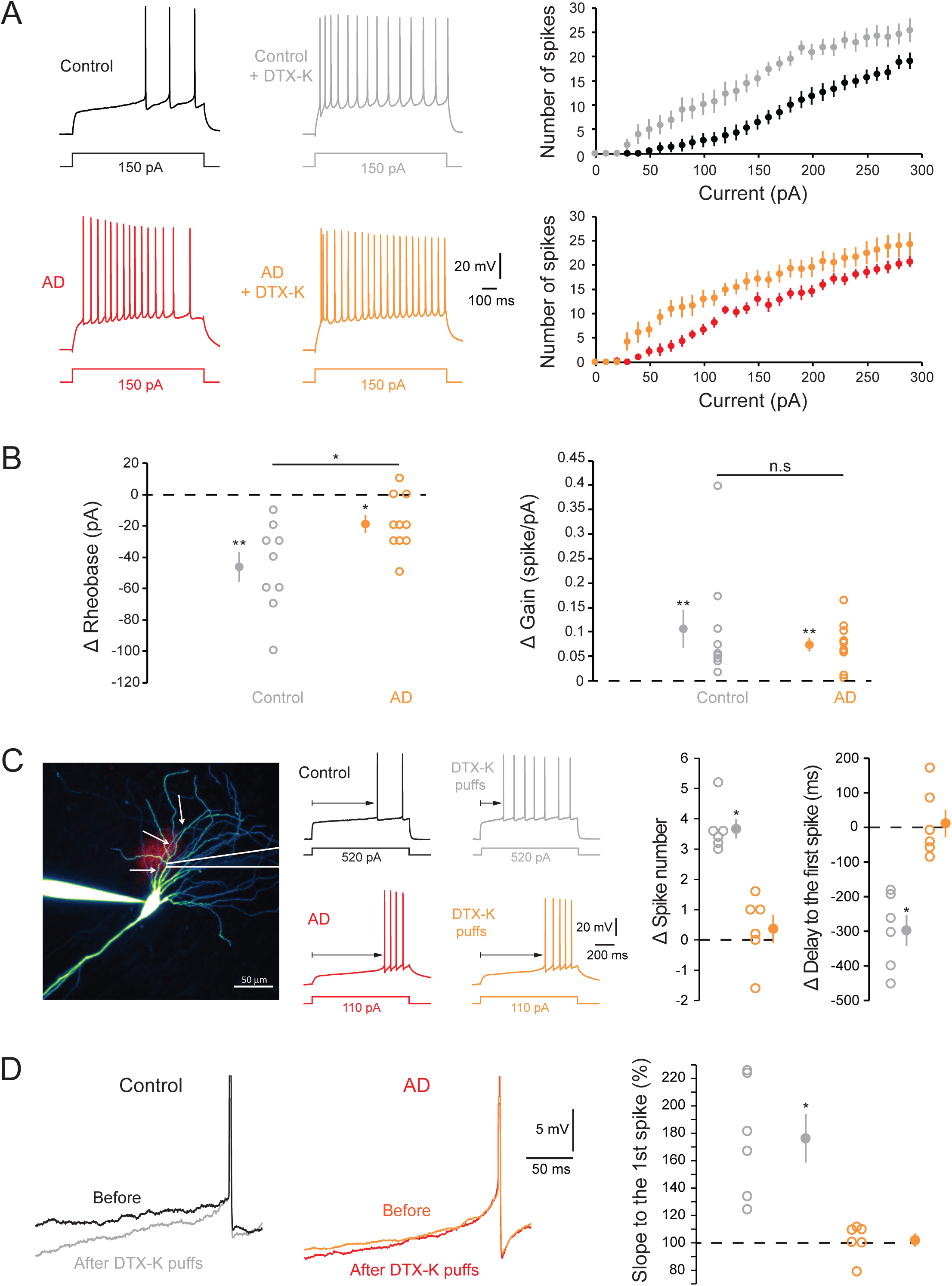
Blockade of axonal K_v_1.1 channels increases IE in AD cultures. A. Increase in intrinsic excitability in control and AD cultures following DTX-K bath application. Left, Example current-clamp traces recorded in CA3 pyramidal neurons in control cultures (black: before DTX-K application; grey: after DTX-K application, n = 9) and in AD cultures (red: before DTX-K application; orange: after DTX-K application, n = 10). Right, Average data across groups. B. Left, effect of DTX-K bath application on rheobase in control cultures (grey) and AD cultures (orange). Note that the rheobase is reduced both in control cultures (Wilcoxon test: p < 0.01) and in AD cultures (Wilcoxon test: p < 0.05) but the reduction is stronger in control cultures (Mann-Whitney test: p < 0.05). Right, effect of DTX-K bath application on input/output curve gain in control cultures (grey) and AD cultures (orange). Note that the gain is increased both in control cultures (Wilcoxon test: p < 0.01) and in AD cultures (Wilcoxon test: p < 0.01) and that this increase is similar in both conditions (Mann-Whitney test: p > 0.1). C-D. DTX-K puffing on axons increase intrinsic excitability in control cultures but not in AD cultures. C. Left, Example of a neuron filled with Alexa 488 while a pipette is puffing DTX-K onto its axon (red spot). The arrows indicate the axon. Middle, Example current-clamp traces recorded in CA3 pyramidal neurons in control cultures (black: before DTX-K puffs; grey: after DTX-K puffs) and in AD cultures (red: before DTX-K puffs; orange: after DTX-K puffs). Right, Statistics of DTX-K axonal puffs effect in control (grey, n = 6) and AD cultures (orange, n = 6). Note that DTX-K puffs increase the number of APs emitted in control cultures but not in AD cultures. Similarly, DTX-K puffs decreases the delay of the first AP emitted in control but not in AD cultures. D. Left, Example of the voltage depolarization slope before the first spike in control cultures (black: before DTX-K puffs; grey: after DTX-K puffs) and in AD cultures (red: before DTX-K puffs; orange: after DTX-K puffs). Right, Statistics of DTX-K axonal puffs effect in control (grey, n = 6) and AD cultures (orange, n = 6). Note that DTX-K puffs increase the voltage depolarization slope before the first spike in control cultures but not in AD cultures.

Immunostaining of K_v_1.1 showed a strong decrease in Kv1.1 staining at the AIS of CA3 pyramidal neurons in AD cultures (**Figure 2)**. Moreover, it has been previously shown that axonal Kv1.1 are a major determinant of CA3 pyramidal neuron intrinsic excitability (*49*). To directly test the implication of axonal Kv1.1 channels in the HP-IE, we locally blocked K_v_1.1 channels by puffing DTX-K on the proximal axon at ∼60 µm from the soma. Interestingly, we found that DTX-K puffing increased IE of CA3 pyramidal neurons in control but not in AD cultures (**Figure 3C)**. In fact, the number of AP elicited by a given current step was increased in control cultures (increase of 3.67 ± 0.32 APs, n = 6; Wilcoxon test: p < 0.05) but not in AD cultures (increase of 0.37 ± 0.46 APs, n = 6; Wilcoxon test: p > 0.1). Similarly, the delay of the first spike was reduced by DTX-K puffing in control cultures (-297.8 ± 44.7 ms, n = 6; Wilcoxon test: p < 0.05) but not in AD cultures (+11.4 ± 40.2 ms, n = 6; Wilcoxon test: p > 0.1). Finally, the depolarizing slope before the first spike was increased by DTX-K puffing in control cultures (176.3 ± 17.6 % of the value before DTX-K puffs, n = 6; Wilcoxon test: p < 0.05) but not in AD cultures (101.9 ± 5 % of the value before DTX-K puffs, n = 6; Wilcoxon test: p > 0.1) (**Figure 3D**). From these results, we conclude that the downregulation of axonal K_v_1.1 channels in AD cultures determines the homeostatic increase in IE in CA3 pyramidal neurons.

### Reduction of Kv1.1 channels increases spike half-width and glutamate release

HP-ST is often attributed to a regulation of the density of postsynaptic receptors. However, it has been previously shown that pharmacological blockade of K_v_1.1 channels increase release probability at CA3-CA3 synapses via broadening of the presynaptic action potential (*45*). Therefore, the increase in ST observed in AD cultures (**Figure 1B**) can be due to presynaptic axonal K_v_1.1 channels down-regulation. Firstly, we calculated the paired pulse-ratio (PPR) of synaptic responses at monosynaptically connected CA3 pairs in control and AD cultures. A significant reduction in the PPR was observed in AD cultures compared to control cultures (control cultures: 99.5 ± 3.3 %, n = 54, AD cultures: 83.01 ± 2.8 %, n = 49, Mann-Whitney test: p<0.001; **Figure 4A**), indicating an increase in glutamate release probability. We conclude that HP-ST observed in AD cultures has a presynaptic component.

**Figure 4:**
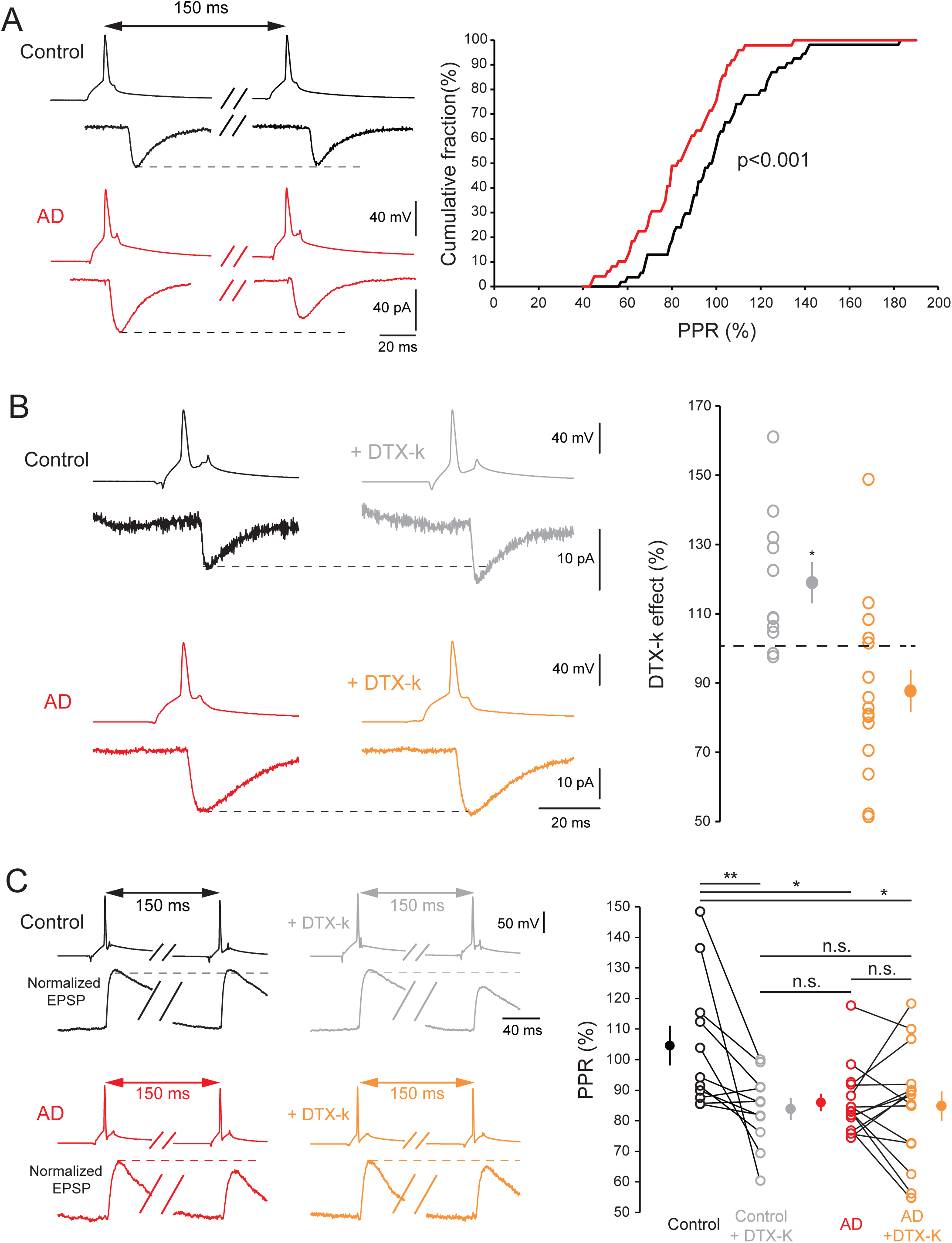
Downregulation K_v_1.1 channels increases glutamate release probability and synaptic strength in AD cultures. A. Smaller Paired Pulse Ratio in AD cultures (red) than in control cultures (black). Left, examples of CA3-CA3 synaptic responses in control and AD cultures (average of 30 traces). Right, cumulative histogram of PPR in control (black) and AD (red) cultures. Note the leftward shift for AD cultures showing a global decrease in PPR. B. K_v_1.1 blockade increases CA3-CA3 synaptic strength in control but not in AD cultures. Left, examples of K_v_1.1 blockade effect on synaptic transmission in control cultures (black: before DTX-K application; grey: after DTX-K application) and AD cultures (red: before DTX-K application; orange: after DTX-K application). Note the increase in synaptic transmission by DTX-K application in control but not in AD cultures. Right, Statistics of DTX-K effect on synaptic strength in control cultures (grey) and AD cultures (orange). C. K_v_1.1 blockade increases release probability at CA3-CA3 synapses in control but not in AD cultures. Left, examples of K_v_1.1 blockade effect on PPR in control cultures (black: before DTX-K application; grey: after DTX-K application) and AD cultures (red: before DTX-K application; orange: after DTX-K application). Note the decrease in PPR by DTX-K application in control but not in AD cultures. Right, Statistics of DTX-K effect on PPR in control and AD cultures (black: control, grey: control + DTX-K, red: AD, orange: AD + DTX-K). Note that the PPR is reduced in control cultures (control v.s control + DTX-K; Wilcoxon test: p < 0.01) but not in AD cultures (AD v.s AD + DTX-K; Wilcoxon test: p > 0.1). Importantly, the PPR is larger in control compared to AD or AD + DTX-K (Mann-Whitney tests: p < 0.05) but is similar between control + DTX-K and AD or AD + DTX-K (Mann-Whitney tests: p > 0.1).

To determine if presynaptic K_v_1.1 channels downregulation could partly explain the increase in release probability, we tested the effect of DTX-K application on synaptic strength in control and AD cultures. DTX-K application increased postsynaptic responses amplitude in control (118.9 ± 6.0 % of the control value, n = 11, Wilcoxon test: p < 0.05) but not in AD cultures (87.4 ± 6.5 % of the control value, n = 15; Wilcoxon test: p > 0.1; **Figure 4B**). Moreover, DTX-K application leads to a decrease in PPR in control (control: 104.6 ± 6.5 %, n = 11; control + DTX-K: 83.9 ± 3.6 %, n = 11; Wilcoxon test: p < 0.01) but not in AD cultures (AD: 86 ± 2.9 %, n = 15; AD + DTX-K: 84.8 ± 4.8 %, n = 15; Wilcoxon test: p < 0.05; **Figure 4C**). Therefore, the increase in ST provoked by K_v_1.1 blockade in control cultures is due to an increase in glutamate release. Importantly, DTX-K application in control cultures was enough to decrease the PPR down to the value observed in AD cultures (Mann-Whitney tests: p > 0.05; **Figure 4C**), showing that K_v_1.1 downregulation in AD cultures is likely to be the cause of the increase in glutamate release. Moreover, in control cultures, we found a strong correlation between the value of the PPR before DTX-K application and the DTX-K induced decrease in the PPR (**Supplementary Figure 3A**), indicating that low release probability in control cultures is partly determined by strong presynaptic K_v_1.1 expression. This correlation was absent in AD cultures (**Supplementary Figure 3B**). We conclude that presynaptic K_v_1.1 downregulation in AD cultures leads to an increase in glutamate release probability at CA3-CA3 synapses and therefore an increase in synaptic strength.

As K_v_1 channels are known to determine synaptic release probability by controlling presynaptic spike width (*26, 34, 45*), we determined spike waveform in control and AD cultures. Firstly, we found that the half-width of the somatic spike was larger in AD cultures than in control cultures (control: 1.33 ± 0.04 ms, n = 80; AD: 1.66 ± 0.06 ms, n = 47; Mann-Whitney test: p < 0.001; **Figure 5A**). Then, we examined axonal spike half-width using voltage imaging and axonal cell-attached recordings (**Figure 5B-D**). Due to the limitation of the dye diffusion, the voltage imaging measurements were done at the AIS (34.2 ± 2.4 µm from the soma; control: n = 7; AD: n = 7) while cell-attached recordings were performed at more distal locations (136.8 ± 17.7 µm from the soma; control: n = 10; AD: n = 7). Importantly, for axonal cell-attached recordings, we measured the peak-to-peak duration of the signal which has been shown to be a good approximation of the spike half-width (*39*) (**Figure 5C**). In fact, we found no statistical difference in axonal spike half-widths measured in voltage-imaging or in cell-attached (**Supp Fig.4**) and therefore we pooled those data. Similar to the somatic spike, we found that the axonal spike was broader in AD cultures than in control cultures (control: 2.12 ± 0.2 ms, n = 17; AD: 2.92 ± 0.27 ms, n = 14; Mann-Whitney test: p < 0.05; **Figure 5D**). Altogether, these results suggest that the chronic activity deprivation by kynurenate leads to a downregulation of axonal K_v_1.1 channels inducing a broadening of axonal AP and the increase in glutamate release probability at CA3-CA3 synapses.

**Figure 5:**
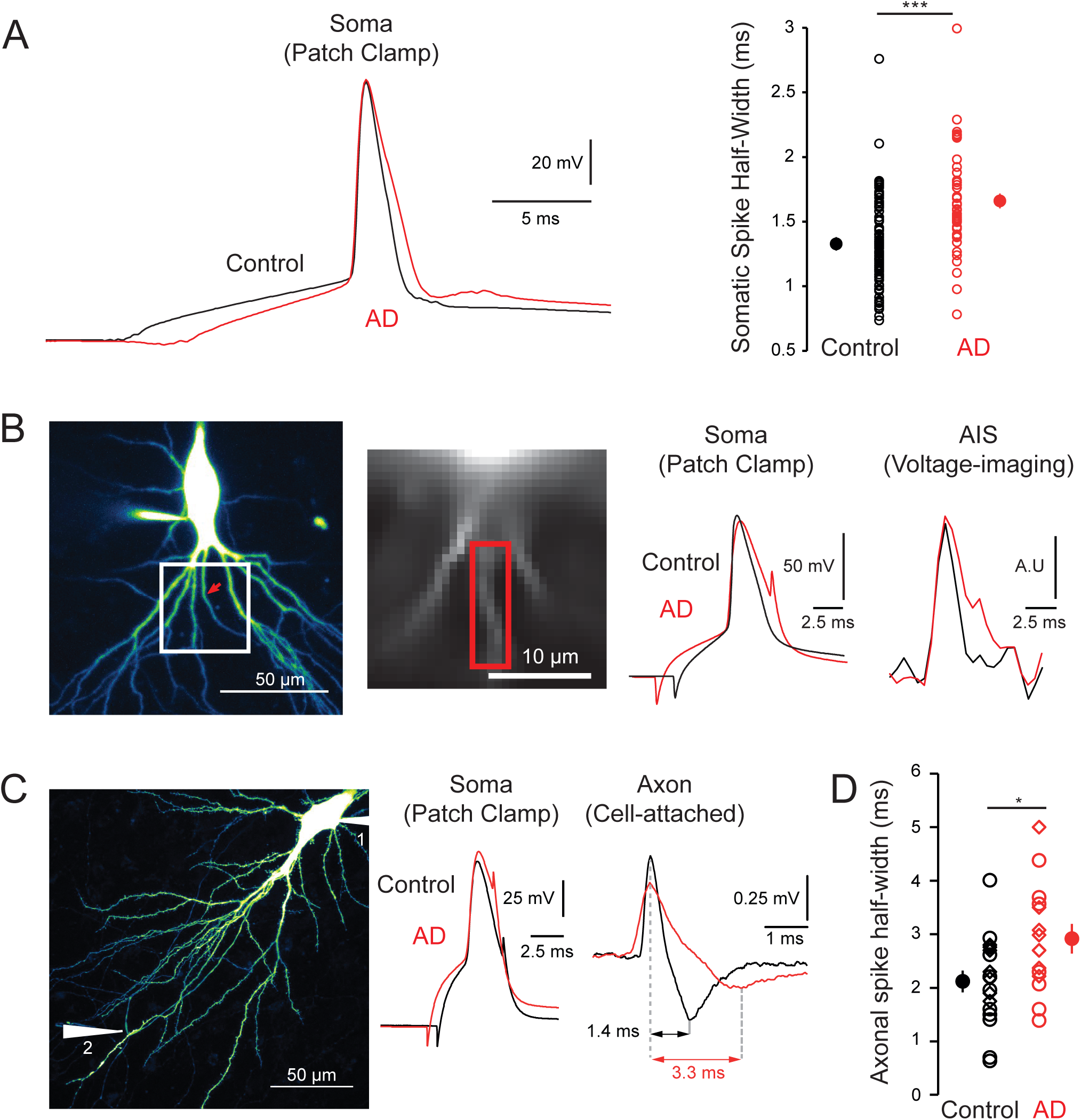
Somatic and axonal action potential are broader in AD cultures. A. Somatic spike is broader in AD cultures (red) compared to control cultures (black). Left, examples of somatic spikes in control and AD cultures. Right, Statistics of somatic spike half-width in control and AD cultures. B-C-D. Axonal spike is broader in AD cultures compared to control cultures. B. Left, Example of a CA3 neuron filled with Alexa 488 and JPW3028. Left image, Morphology of the neuron obtained following excitation of Alexa 488 (red arrow: axon). Right image, Fluorescence image obtained following excitation of JPW3028 (red rectangle: ROI used to measure voltage-imaging signal). Right, Example of spikes recorded in the soma with electrophysiology (left) and in the AIS with voltage imaging (right) (control: black; AD: red). Note that the axonal spike is broader in AD cultures. C. Left, Example of a CA3 neuron filled with Alexa 488. The axon collateral (white arrow number 2) is identified and recorded in a cell-attached configuration. Right, Examples of spikes recorded in the soma in whole-cell mode and in the axon in cell-attached mode (control: black; AD: red). Note that the peak-to-peak duration of the axonal signal is larger in AD cultures. D. Statistics of axonal spike half-width in control and AD cultures (empty rhombi: individual voltage-imaging data points, empty dots: individual cell-attached data points, full dots: means).

### Increase in spontaneous mEPSCs but not evoked mEPSCs in AD neurons

To test whether the increase in synaptic strength in AD cultures goes with an increase in postsynaptic receptors density we recorded spontaneous miniatures EPSCs (mEPSCs) in CA3 neurons of control and AD organotypic cultures while blocking spiking with 1 µM TTX (tetrodotoxin), inhibitory transmission with 100 nM PTX (picrotoxin) and mossy fiber synapses with 1 µM DCG IV (*50*). An increase in amplitude of spontaneous mEPSCs was found in AD cultures (136.4 %; control: -10.11 ± 0.61 pA, n = 16; AD cultures: -13.8 ± 0.84 pA, n = 19; Mann-Whitney test: p < 0.05; **Figure 6A**), indicating that postsynaptic AMPAR density was increased in AD cultures. Furthermore, spontaneous mEPSCs frequency was higher in AD cultures compared to control cultures (**Figure 6A**), probably due to an increase in presynaptic vesicles release probability.

**Figure 6:**
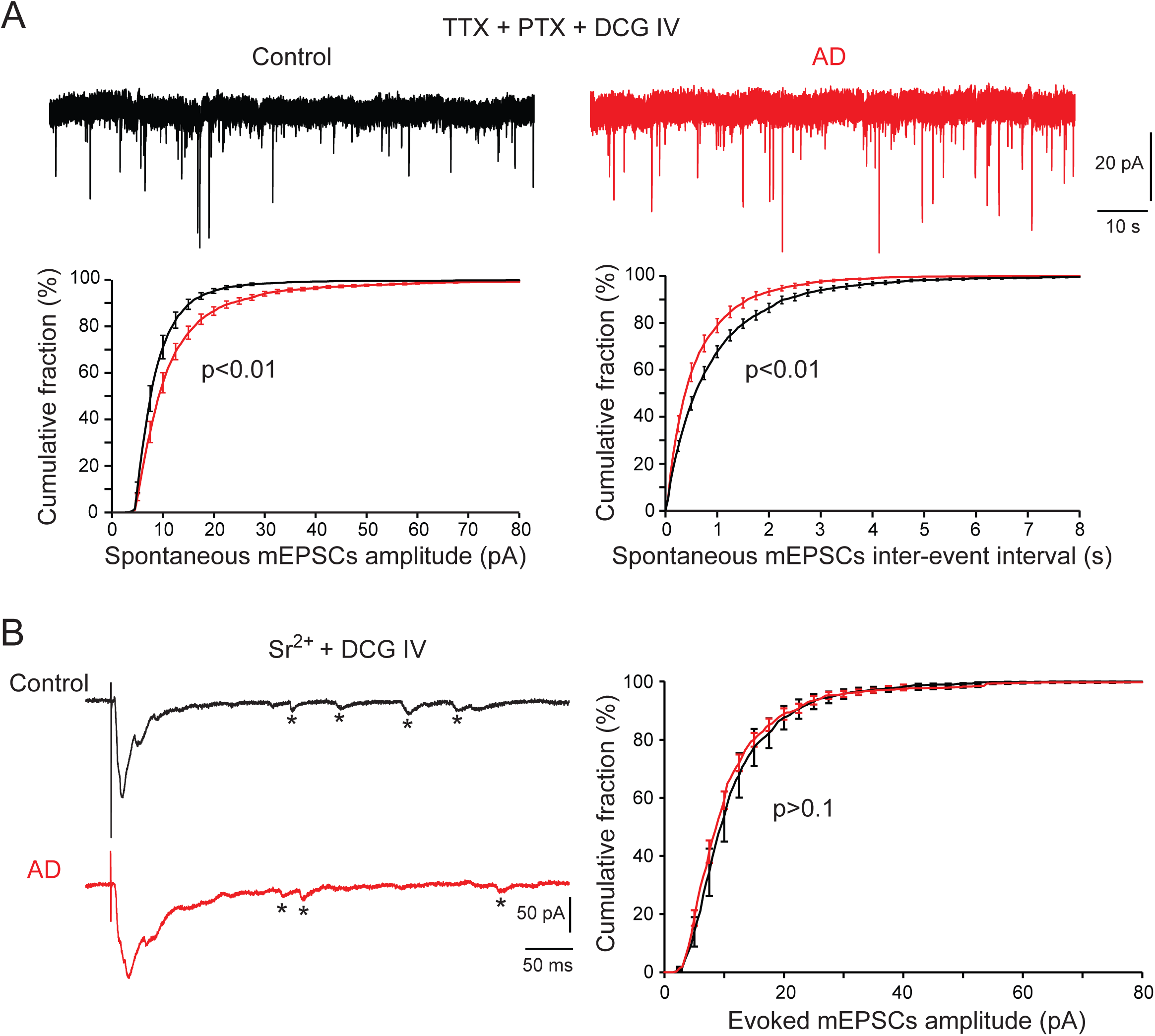
Increase in *spontaneous* but not *evoked* mEPSCs amplitude in AD cultures. A. Spontaneous mEPSCs display increased amplitude and frequency in AD cultures. Up, examples of 1-minute recordings of spontaneous mEPSCs (recorded in the presence of TTX, PTX and DCG IV) in control (black) and AD (red) cultures. Bottom left, cumulative histogram of spontaneous mEPSCs amplitude in control (black) and AD (red) cultures. Note the rightward shift for AD cultures showing an increase in amplitude. Bottom right, cumulative histogram of spontaneous mEPSCs inter-event interval in control (black) and AD (red) cultures. Note the leftward shift for AD cultures showing an increase in frequency. B. Evoked mEPSCs show no difference in amplitude in control and AD cultures. Left, examples of evoked mEPSCs (recorded 500 ms following extracellular stimulation in the presence of Sr^2+^ and DCG IV) in control (black) and AD (red) cultures. Right, cumulative histogram of evoked mEPSCs amplitude in control (black) and AD (red) cultures.

However, previous studies showed that the pool of vesicles spontaneously released is different from the one released following spiking (*51–54*). Some studies even showed that these two vesicles pools activate different pools of postsynaptic receptors (*55, 56*). Therefore, the increase in evoked synaptic transmission we measured in AD cultures (**Figure 1A**) could be independent of the increase in mEPSC amplitude and frequency (**Figure 6A**). To answer this question, we decided to record the amplitude of mEPSCs directly evoked by presynaptic spiking. To do so, we evoked monosynaptic EPSCs by extracellular stimulation in CA3 pyramidal layer while substituting extracellular Ca^2+^ with Sr^2+^ to induce desynchronization of the presynaptic release (*57*). In order to keep the firing, no TTX was present in the bath. We blocked mossy fiber transmission with 1 µM DCG IV but GABAergic inhibition was left intact to avoid epileptiform activity. Importantly, to avoid the detection of GABAergic events we used a low-chloride intracellular solution, and we recorded the neurons at the Cl^-^ reversal potential (-70 mV in this case). We measured the amplitude of the desynchronized synaptic events during the 500 ms following the extracellular stimulation (stars in **Figure 6B**). These events are due to the desynchronised fusion of single vesicles following the invasion of the presynaptic terminal by an action potential (*57*). We called them evoked mEPSCs. Interestingly, we found no difference in evoked mEPSCs amplitude in control and AD cultures (control: -12.08 ± 1.42 pA, n = 11; AD cultures: -12.09 ± 0.58 pA, n = 12; Mann-Whitney test: p > 0.1; **Figure 6B**). Therefore, we found an increase in spontaneous mEPSCs amplitude but no difference in evoked mEPSCs amplitude in AD cultures. This raises the possibility that the increase in postsynaptic receptors density does not participate in the HP-ST found in AD cultures.

### Depolarization induced Analog-Digital Facilitation is absent in AD cultures

In CA3 pyramidal neurons, a 10 seconds somatic subthreshold depolarization inactivates axonal K_v_1.1 channels leading to a broadening of the axonal spike and an increase in synaptic release (*31, 45*). This facilitation is named d-ADF for depolarization induced Analog-Digital Facilitation (*46, 58*). As axonal K_v_1.1 channels are down-regulated in AD cultures, we could expect a disappearance of d-ADF. Firstly, we examined the depolarization-induced broadening of axonal spike using voltage imaging and cell attached axonal recordings (**Figure 7A-B**). Somatic depolarization from -80 mV to -50 mV for 10 s leaded to a 30.6 % broadening of axonal spike in control cultures (-80 mV: 2.42 ± 0.18 ms, -50 mV: 3.17 ± 0.23 ms, n = 13, Wilcoxon test: p < 0.01; **Figure 7C**) but had no effect in AD cultures (-80 mV: 2.92 ± 0.28 ms, -50 mV: 2.88 ± 0.27 ms, n = 14, Wilcoxon test: p > 0.1; **Figure 7C**). Therefore, the axonal spike broadening following subthreshold somatic depolarization is absent in AD cultures.

**Figure 7:**
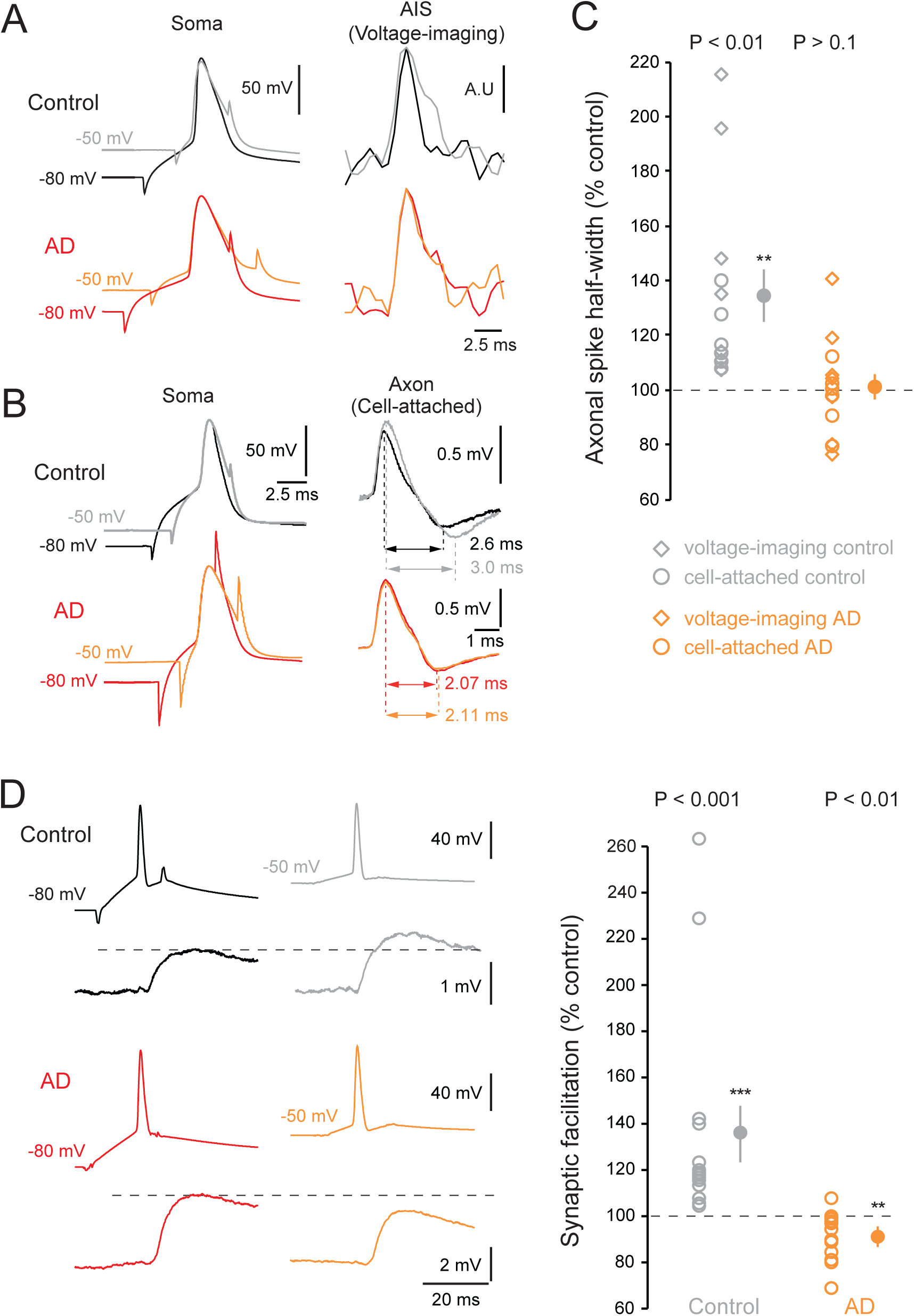
Loss of depolarization-induced axonal spike broadening and synaptic facilitation in AD cultures. A-B-C. Disappearance of depolarization induced axonal spike broadening in AD cultures. A. Examples of spikes recorded in the soma with electrophysiology and in the AIS with voltage imaging. Note the broadening of axonal spike at depolarized membrane potential in control cultures (black: hyperpolarized membrane potential; grey: depolarized membrane potential) but not AD cultures (red: hyperpolarized membrane potential; orange: depolarized membrane potential). B. Examples of spikes recorded in the soma in whole-cell mode (left) and in the axon in cell-attached mode (right). Note the increase in peak-to-peak duration of the axonal spike at depolarized membrane potential in control cultures (black: hyperpolarized membrane potential; grey: depolarized membrane potential) but not AD cultures (red: hyperpolarized membrane potential; orange: depolarized membrane potential). C. Statistics of subthreshold depolarization effect on axonal spike width in control cultures (grey) and AD cultures (orange) (empty rhombi: individual voltage-imaging data points, empty dots: individual cell-attached data points, full dots: means). D. Reversion of depolarization induced Analog Digital facilitation (d-ADF) in AD cultures. Left up, example of CA3-CA3 connection showing presynaptic depolarization induced facilitation in a control culture (black: hyperpolarized presynaptic membrane potential; grey: depolarized presynaptic membrane potential). Left down, CA3-CA3 connection shows a presynaptic depolarization induced depression in an AD culture (red: hyperpolarized presynaptic membrane potential; orange: depolarized presynaptic membrane potential). Right, Statistics of presynaptic depolarization effect on synaptic strength in control cultures (grey) and AD cultures (orange).

Next, we tested the increase in glutamate release following subthreshold depolarization by recordings of pairs of monosynaptically connected CA3 cells. We depolarized the presynaptic cell resting membrane potential to -50 mV for 10 s. This protocol induced an increase in the postsynaptic response amplitude in control cultures (133.9 ± 12.0 %, n = 15; Wilcoxon test: p<0.001; **Figure 7C**), showing that d-ADF is present at CA3-CA3 synapses. However, in AD cultures, presynaptic depolarization led to a decrease in synaptic transmission (89.7 ± 2.9 % of the control value, n = 13, Wilcoxon test: p < 0.01; **Figure 7C**). Thus, d-ADF is reversed in AD cultures. Interestingly, this result is consistent with a previously published result showing that pharmacological blockade of K_v_1.1 channels induces a reversion of d-ADF in control cultures (see Figure 2 in (*45*)). Therefore, the reversion of d-ADF in AD cultures is likely to be a consequence of K_v_1.1 downregulation, probably due to the unravelling of sodium channels inactivation following presynaptic depolarization (*59–62*). Altogether, these results show that K_v_1.1 downregulation following chronic activity blockade induces the disappearance of K_v_1.1-dependent increase in axonal spike duration and synaptic transmission induced by somatic depolarization.

## Discussion

Our study demonstrates a synergy between homeostatic plasticity of intrinsic excitability (HP-IE) and homeostatic plasticity of synaptic transmission (HP-ST) in CA3 network following activity deprivation. This synergy could be efficiently explained by the down-regulation of a single molecular actor: the axonal K_v_1.1 channels. In fact, K_v_1.1 channels located in the AIS and proximal axon control IE in CA3 pyramidal cells (*3, 49*) whereas K_v_1.1 channels located in presynaptic terminals control presynaptic spike duration and synaptic release (*31, 45, 46, 63*). Thus, the downregulation of axonal K_v_1.1 channels following chronic AD leads to the increase in intrinsic excitability of CA3 neurons and synaptic strength at CA3-CA3 contacts. Furthermore, I_D_ current downregulation reverses d-ADF. In fact, a presynaptic subthreshold depolarization *increases* synaptic release in control cultures but *decreases* it in AD cultures.

### Synergistic homeostatic regulation of IE and ST following activity deprivation

Various studies showed a synergy of HP-IE and HP-ST in neuronal networks (*17, 64, 65*). Usually, this synergy is explained by common molecular pathways leading to regulation of different molecular effectors. For example, in dissociated cortical cultures, chronic activity deprivation leads to a diminution of BDNF exocytosis entailing both HP-ST through regulation of AMPARs and of HP-IE through regulation of sodium and potassium voltage-gated channels (*4, 5, 66*). In CA3 network, chronic activity deprivation has been shown independently to induce HP-IE (*3*) and HP-ST (*20*). Here, we show that these two forms of plasticity act synergistically (**Figure 1**) and are determined partly by the same molecular cause: the downregulation of axonal K_v_1.1 channels. In fact, pharmacological blockade of K_v_1.1 channels increases IE of CA3 neurons to a larger extent in control cultures than in AD cultures (**Figure 3A-B**). Moreover, specific blockade of axonal Kv1.1 channels increases IE of CA3 neurons in control cultures but not in AD cultures (**Figure 3C-D**). Similarly, K_v_1.1 blockade increases ST at CA3-CA3 synapses in control cultures but has no effect in AD cultures (**Figure 4B**). In addition, synaptic connections in AD cultures display smaller PPR than control cultures, indicating that the increase in ST in AD cultures is partly due to an increase in release probability (**Figure 4A**). Interestingly, K_v_1.1 blockade decreases the PPR observed in control cultures down to the value observed in AD cultures (**Figure 4C**). This suggests that the increase in glutamate release probability at CA3-CA3 synapses in AD cultures is due to K_v_1.1 downregulation. Moreover, both somatic and axonal spikes have been found to be broader in AD cultures compared to control cultures. Therefore, it is likely that the downregulation of K_v_1.1 channels located in the AIS and proximal axon accounts for the increase in IE while the downregulation of K_v_1.1 channels in presynaptic terminals accounts for the increase in glutamate release via broadening of the presynaptic spike. We thus describe here a HP-ST via presynaptic spike waveform modulation, a phenomenon that has been described in neocortical dissociated cultures and in batracian neuro-muscular junction (*36, 38*).

### Kv1.1 downregulation is not the only determinant of homeostatic plasticity

In this study, we showed that K_v_1.1 downregulation induces both HP-IE and HP-ST following activity deprivation. However, it is likely that K_v_1.1 downregulation is accompanied by other mechanisms of homeostatic plasticity. First, we found that the decrease of Kv1.1 expression at the AIS in AD cultures went with a 40% increase of AIS length in CA3 pyramidal neurons (**Figure 2**). Increase of AIS length following activity deprivation has been described as a main determinant of HP-IE in several studies (*48, 67–71*). Therefore, it is likely that the increase in AIS length we found is involved in the intrinsic excitability increase of CA3 pyramidal neurons in AD cultures. Second, homeostatic elevation of excitatory synaptic transmission is often due to upregulation of postsynaptic AMPA receptors and/or upregulation of vesicles release probability. To test this hypothesis, we recorded *spontaneous* mEPSCs in presence of sodium channel blocker. These *spontaneous* mEPSCs are due to spike-independent fusion of single glutamatergic vesicles. We found that *spontaneous* mEPSCs amplitude and frequency were increased in AD cultures (**Figure 6A**). Therefore, it is probable that the postsynaptic glutamate receptors and the presynaptic vesicles machinery are also regulated to produce an increase in synaptic strength in AD cultures. However, it has been showed that spontaneous and evoked synaptic transmissions rely on different pools of vesicles and can activate different pools of postsynaptic receptors (*51, 52, 55, 56*). Therefore, we decided to record spike-induced *evoked* mEPSCs. To do so, we substituted Ca^2+^ by Sr^2+^ in extracellular solution, which provokes the desynchronization of vesicles fusion when a spike invades the presynaptic terminal, allowing the measurement of single vesicle fusion events, *i.e evoked* mEPSCs, in the postsynaptic cell. We found no change in *evoked* mEPSCs amplitude in AD cultures (**Figure 6B**). This raises the possibility that the increase in spontaneous synaptic transmission is due to an increase in postsynaptic receptors density while the increase in evoked synaptic transmission is due to presynaptic K_v_1.1 downregulation. Finally, whether regulation of other type of voltage-gated channels (such as up-regulation of presynaptic Nav channels) participate to the enhanced synaptic release at AD synapses cannot be totally discarded.

### Comparison with homeostatic plasticity induced by TTX

Mitra and colleagues found in 2012 that the increase in synaptic transmission between CA3 pyramidal cells following chronic activity blockade goes along with a strong reduction of the connection probability between CA3 neurons (from ∼45% in control cultures to ∼20% in AD cultures). They conclude that the decrease in connectivity avoids the over-excitation of CA3 network despite the increase in synaptic transmission. Here, we found a much smaller and non-significant decrease in connectivity following activity deprivation (48% in control cultures, 39% in AD cultures). This discrepancy may arise from the difference in the drug used to deprive activity in the two studies. Mitra and colleague used TTX, which blocks spiking, whereas we used kynurenate which blocks excitatory synaptic transmission. In their study, Mitra and colleagues show that the decrease in connectivity is due to an increase in the binding between CDK5 and p25/p35, which is known to silence synapses. Interestingly, they do not observe this upregulation of CDK5 after activity blockade with a blocker of excitatory transmission, NBQX. Thus, HP-ST following activity deprivations by spiking blockade (TTX) or excitatory transmission blockade (kynurenate or NBQX) seem to involve different molecular actors (*12, 15, 72*).

### Over-excitation dampening through reversion of d-ADF

In pairs of connected CA3 pyramidal neurons, a long subthreshold depolarization preceding the spike (5-10 s at -50 mV) inactivates axonal K_v_1.1 channels, broadens presynaptic spike and increases synaptic transmission (which produces d-ADF). Thus, a local excitation of the network will be amplified through this phenomenon. In activity deprived (AD) cultures, we showed that K_v_1.1 channels are downregulated and d-ADF is reversed. In this case, we suppose that the absence of K_v_1.1 channels unravels the Na_v_ channels inactivation by subthreshold depolarization leading to decrease in presynaptic spike amplitude and synaptic strength (*59, 61, 62*). This hypothesis will need more experiments to be confirmed. Interestingly, the reversion of d-ADF will tend to dampen over-excitation in CA3 network as a global depolarization will diminish synaptic transmission. Thus, this phenomenon could be a mechanism avoiding over-excitation in AD cultures despite the global increase in intrinsic excitability and synaptic strength. In that point of view, downregulation of K_v_1.1 channels following activity deprivation could allow homeostatic increase in network activity while avoiding pathological states.

### Voltage-gated channels as regulators of synaptic transmission

Classically, the role of voltage gated channels is reduced to determination of neuronal intrinsic excitability. However, numerous studies showed that short-term inactivation of potassium voltage gated channels by resting membrane potential depolarization broadens the action potential and therefore increases synaptic release (*25, 26, 28, 34, 35, 45*). Here, we show that long term downregulation of K_v_1.1 channels entails a broadening of presynaptic spikes and increase in synaptic transmission. Thus, neuronal networks can present long term plasticity of action potential waveform. Moreover, sodium channels availability determines also spike waveform and synaptic transmission (*27, 46, 62, 73*). Therefore, we could expect that a HP-IE via Na_v_ regulation (*2, 5, 6, 8*) may also impact synaptic transmission. In conclusion, regulation of axonal voltage-gated channels could allow neurons to modulate efficiently intrinsic excitability and synaptic transmission as a whole.

## Methods

### Organotypic hippocampal slice cultures

Our experiments were conducted according to the European and Institutional guidelines (Council Directive 86/609/EEC and French National Research Council and approved by the local health authority (Préfecture des Bouches-du-Rhône, Marseille)). Hippocampal slice cultures were prepared using an interface technique (*74, 75*). Briefly, postnatal day 5–7 Wistar rats were deeply anesthetized by intraperitoneal injection of chloral hydrate, the brain was removed, and each hippocampus was individually dissected. Hippocampal slices (300 µm) were placed on 20 mm latex membranes (Millicell) inserted into 35 mm Petri dishes containing 1 ml of culture medium and maintained for up to 12 d in an incubator at 34°C, 95% O_2_–5% CO _2_. The culture medium contained (in ml) 25 MEM, 1.25 HBSS, 12.5 horse serum, 0.5 penicillin/streptomycin, 0.8 glucose (1 M), 0.1 ascorbic acid (1 mg/ml), 0.4 HEPES (1 M), 0.5 B27, and 8.95 sterile H_2_O. To arrest glial proliferation, 5 µM Ara-C was added for one night to the culture medium after 3 days *in vitro* (DIV).

### Solutions and pharmacology

The control perfusion solution contained (in mM) 125 NaCl, 26 NaHCO_3_, 3 CaCl_2_, 2.5 KCl, 2 MgCl_2_, 0.8 NaH_2_PO_4_, and 10 D-glucose and was continuously equilibrated with 95% O_2_–5% CO_2_. Patch pipettes (6 –9 MΩ) were filled with a solution containing (in mM) 120 K-gluconate, 20 KCl, 10 HEPES, 0.5 EGTA, 2 Na_2_ATP, 0.3 NaGTP, and 2 MgCl_2_, pH 7.4. All drugs were applied via the bath solution. D-type currents (K_v_1 channels) were blocked by bath application of DTX-K (50 –100 nM; Sigma), a blocker of K_v_1.1 channels. To chronically block excitatory transmission, 2 mM Kynurenate was added to the culture medium at 2 or 3 days before recordings; recordings of treated cultures were performed 7–12 DIV.

### Paired recordings and analysis

Dual recordings from pairs of neurons were obtained as previously described (*74*). The external saline contained (in mM): 125 NaCl, 26 NaHCO_3_, 3 CaCl_2_, 2.5 KCl, 2 MgCl_2_, 0.8 NaH_2_PO_4_ and 10 D-glucose, and was equilibrated with 95% O_2_–5% CO_2_. Patch pipettes (5–10 MO) were pulled from borosilicate glass and filled with an intracellular solution containing (mM): 120 K gluconate, 20 KCl, 10 HEPES, 0.5 EGTA, 2 MgCl_2_, 2 Na_2_ATP and 0.3 NaGTP (pH 7.4). Recordings were performed at 30°C in a temperature-controlled recording chamber (Luigs & Neumann, Ratingen, Germany). Usually, the presynaptic neuron was recorded in current clamp and the postsynaptic cell in voltage clamp. Both pre- and postsynaptic cells were held at their resting membrane potential (approximately -65 mV). Presynaptic APs were generated by injecting brief (5 ms) depolarizing pulses of current at a frequency of 0.1 Hz. PPR was assessed with two presynaptic stimulations (150-ms interval). Voltage and current signals were low-pass-filtered (3 kHz), and sequences (200–500 ms) were acquired at 10–20 kHz with pClamp (Axon Instruments, Molecular Devices) version 10. Electrophysiological signals were analysed with ClampFit (Axon Instruments) and custom-made softwares written in LabView (National Instruments). Postsynaptic responses could be averaged following alignment of the presynaptic APs using automatic peak detection. The presence or absence of a synaptic connection was determined on the basis of averages of 20 individual traces.

### mEPSCs recordings and analysis

In *spontaneous* mEPSCs experiments, inhibitory synaptic transmission, Nav channels and mossy fiber synapses were blocked using respectively 100 nM picrotoxin (PTX), 1 µM TTX (tetrodotoxin) and 1 µM DCG-IV (mGluR2 agonist).

In *evoked* mEPSCs experiments, only mossy fiber synapses were blocked using 1 µM DCG-IV. To induce desynchronization of evoked synaptic transmission Ca^2+^ was replaced by Sr^2+^ in extracellular solution. To avoid the detection of GABAergic events in this experiment we used a low-chloride intracellular solution (in mM: 150 K gluconate, 4.6 KCl, 10 HEPES, 0.5 EGTA, 2 MgCl_2_, 2 Na_2_ATP and 0.3 NaGTP, 10 phosphocreatine) and we recorded the neurons at the Cl^-^ reversal potential (-70 mV in this case).

Spontaneous and evoked mEPSCs were selected by hand using ad hoc analysis software written in LabView (National Instruments). Then the traces were exported in ClampFit (Axon Instruments) to analyse the amplitude and inter-event intervals.

### Axonal cell-attached recordings

Simultaneous recordings from the soma in whole-cell configuration and the axon in cell-attached configuration were obtained from CA3 pyramidal neurons. Briefly, CA3 pyramidal cells were recorded with 50 µM Alexa 488 (Invitrogen) in the recording solution. After whole cell access of the somatic compartment, we waited 10-15 minutes to allow the dye to strongly stain the axonal compartment and visualized the cell at 488 nm with a LSM-710 confocal microscope (Zeiss). The axon was identified by visual clues: a thin, aspiny process emerging from the soma or a proximal dendrite.

We approached a secondary pipette containing extracellular solution to the desired axonal target and a brief suction was applied to the pipette. Under these conditions, the spike measured in the axon had a positive polarity (amplitude 0.3-1.2 mV) when recorded in current clamp.

### Puff applications

CA3 neurons were filled with Alexa 488 50 µM (Invitrogen) to visualize neuronal morphology. Neurons were recorded for ∼10 minutes before imaging with a LSM 710 confocal system (Zeiss). Alexa 488 was excited with laser source at 488 nm. The axon was identified by visual clues: a thin, aspiny process emerging from the soma or a proximal dendrite.

Localized puffs of K_v_1.1 channel blocker were applied on the AIS of the recorded neuron with a patch-pipette filled with a solution containing the extracellular saline with 50 µM Alexa 594 (Invitrogen) and 100 nM DTX-K (Sigma). Alexa 594 was excited with laser source at 543 nm. Short puffs (20–50 ms, 5 psi) were emitted every 10 s by a Toohey-Spritzer pressure system (Toohey Company).

### Voltage imaging of the axon

CA3 pyramidal neurons were imaged with JPW3028 (250 µM) following a protocol detailed by (*44*). JPW3028 was a gift of Dr L. M. Loew (University of Connecticut Health Center, Farmington, CT, USA). In practice, the tip of the electrode was pre-filled with Alexa-488 (50 µM) to visualize the morphology of the neuron and back-filled with the voltage sensitive dye JPW-3028 (250 µM). After 5 min of whole-cell recording, Alexa diffused in the neuron and the neuronal morphology was acquired with the LSM- 710 confocal microscope. If the neuron displayed a clearly visible axon and axon initial segment (AIS), the protocol was continued (otherwise the cell was discarded). The AIS was defined as the proximal segment of the axon extending up to 60 µm from the axon hillock in CA3 pyramidal neurons (*76*). The cell was recorded in whole-cell mode for at least 25 min to obtain sufficient diffusion of the dye into the axon, and the imaging protocol was then started. The cell was illuminated in wide field with a 525-nm LED system (CoolLed; Roper Scientific/Photometrics, Tucson, AZ, USA) via a 60 × 1.0 N.A. water-immersion objective (Zeiss). Collected fluorescence was long-pass filtered at 610 nm and projected onto a 128 × 128-pixel high-speed EMCCD camera (Evolve 128; Photometrics) with a maximum frame rate of 7 kHz. In a typical trial, we induced APs in the soma by injecting a 5- to 10-ms current pulse, synchronized with LED illumination and camera acquisition at 500 Hz for 40 ms. To improve the signal-to-noise ratio, we acquired 50–100 sweeps sequentially and reconstructed the signal at 1-2 kHz by a shift and mean algorithm (*77*). All these operations were performed using ad hoc analysis software written in LabView (National Instruments).

### Immunohistochemistry

Control and activity deprived hippocampal organotypic cultures were fixed in 4% paraformaldehyde for 30 minutes and washed in PBS. Immunodetection was done in free floating sections. Brain slices were treated with 50 mM NH_4_Cl for 30 minutes and incubated in blocking buffer (10% goat serum, 0.1 % Triton X-100 in PBS) for 2 hours to avoid nonspecific binding. Slices were incubated overnight at 4° C with the primary antibodies diluted in incubation buffer (PBS, 1% goat serum and 0.1% Triton X-100). The primary antibodies used were: mouse IgG2b anti-K_v_1.1 (1:100, Neuromab K36/15) and mouse IgG2a anti-ankyrinG (1:100, Neuromab N106/36). After extensive washing in incubation buffer, the secondary antibodies, Alexa-568 goat anti-mouse IgG2a (1:500) and Alexa-488 goat anti-mouse IgG2b (1:500) were incubated 2 hours at room temperature and then washed. Bisbenzimide (1:2500) was added for 3 minutes to stain nuclei and identify hippocampal formation domains. After staining, the coverslips were mounted with Fluoromount-G (Southern-Biotech) and the images were acquired on a confocal microscope (Leica SP5) using the same parameters to compare intensities between control and experimental conditions. Images were prepared for presentation using the Adobe CS4 software.

### Immunochemistry image analysis and statistics

Kv1.1 fluorescence intensity was analyzed using ImageJ/Fiji software in AIS and neuronal somas from 11 control and 11 treated brain slices obtained from 3 independent experiments. For every AIS, a 5 pixels’ line was drawn along the Kv1.1 staining to obtain fluorescence intensity. Kv1.1 fluorescence integrated density was obtained from at least 10 AISs in each slice. Background integrated density was measured in each slice and corrected total fluorescence (CTF) for each AIS was calculated using the following formula: CTF = Kv1.1 Integrated density – (Kv1.1 Area x Mean fluorescence of background readings). Neuronal somas Kv1.1 CTF was calculated for each slice by drawing a polygon that contains neuronal somas based on nuclei staining. All data were also normalized to the mean of Kv1.1 signal in AISs or somas of control slices. AIS length was measured using ankyrinG staining as previously described (*78*). A line was traced from soma along ankyrinG staining and starting and end position were defined as the position where ankyrinG intensity was 33% of maximum intensity measured in each AIS.

### Statistical tests

The results are presented as the mean mean ± s.e.m. Statistical differences between experimental conditions were calculated not assuming Gaussian distribution or equal variances, using Mann–Whitney U-test or Wilcoxon rank-signed test. For analysis of the connectivity, we used a χ^2^ test.

## Acknowledgements

This work was supported by INSERM, CNRS (to DD), Ecole Normal Supérieure (doctoral grant to MZ), Agence Nationale de la Recherche (REPREK ANR-11-BSV16-016-01 to DD; LoGIK ANR-17-CE16-022), Fondation pour la Recherche Médicale (doctoral grant to MZ FDT-2015-0532147; DVS-2013-1228768 to DD) and Ministerio de Ciencia y Universidades (RTI2018-095156-B-100 to JJG).

## Authors contributions

M.Z. and D.D. conceived the study; M.Z., S.R., L.F.-M., N.B.-G. and A.B. acquired electrophysiological data; M.Z. and S.R. analyzed electrophysiological data; S.R., performed axonal cell-attached recordings and voltage imaging; M-J.B. and J.J.G. performed and analyzed immunohistochemistry; M.Z., S.R., and D.D. wrote the manuscript. All authors approved the final version of this manuscript.

---

**Supplementary Figure 1:**
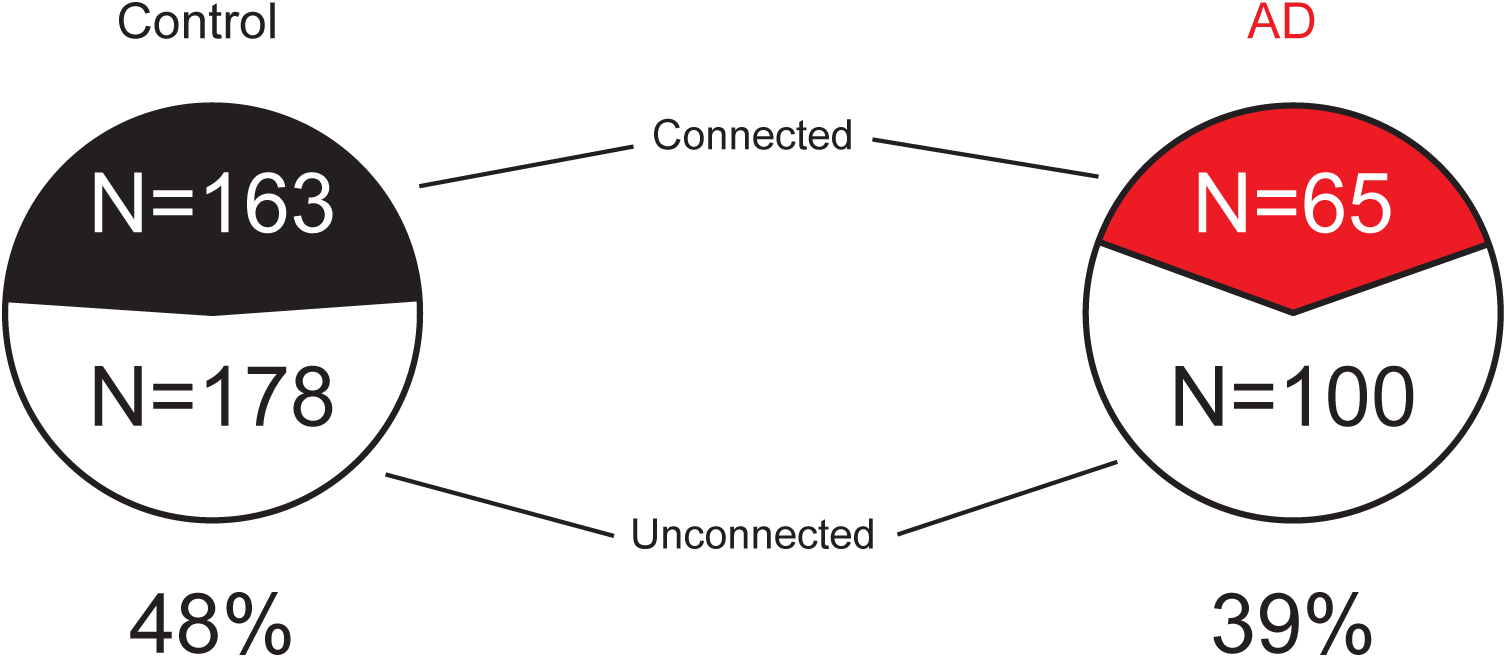
Effect of chronic activity deprivation on connectivity between CA3 neurons. Smaller connectivity between CA3 neurons in activity deprived cultures than in control cultures. Left, connectivity in control cultures (white: unconnected pairs, black: connected pairs). Right, connectivity in AD cultures (white: unconnected pairs, red: connected pairs).

**Supplementary Figure 2:**
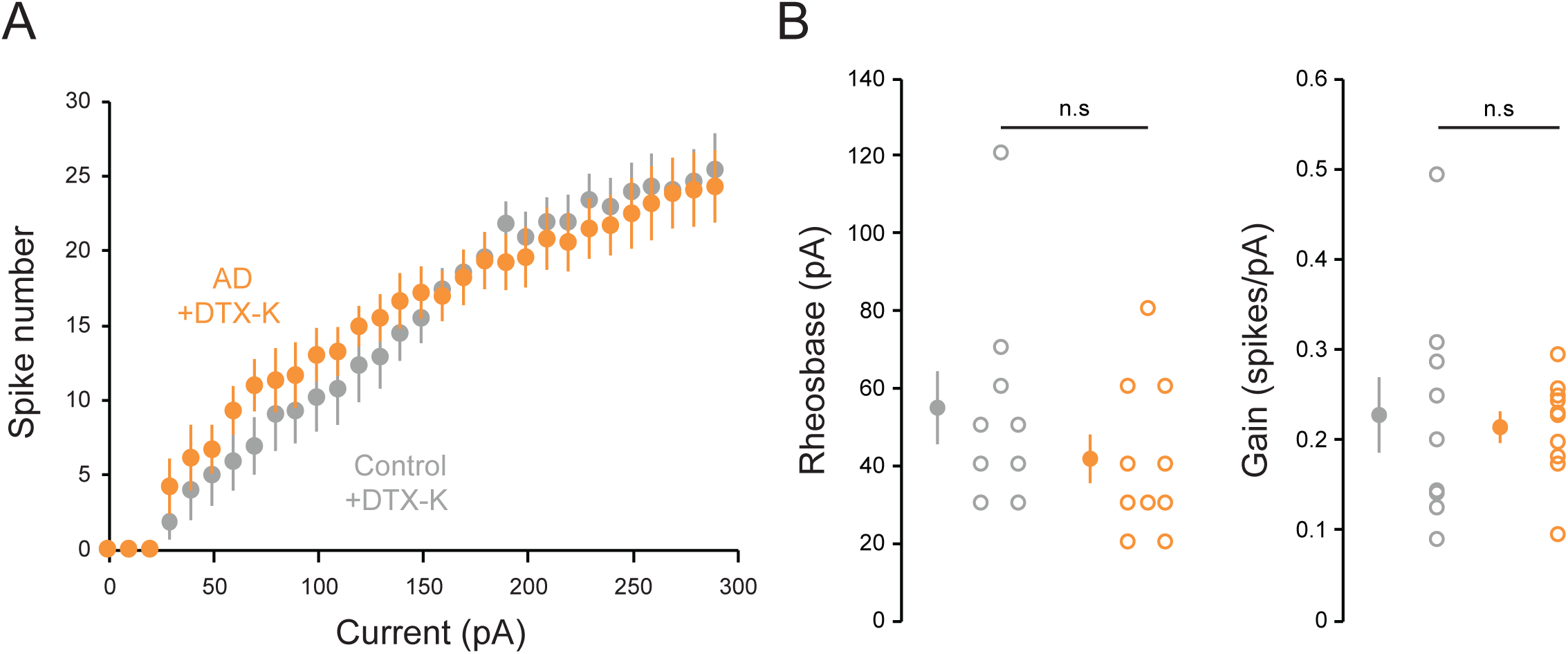
Similar IE in control and AD cultures after DTX-K application. Left, Input/output curves of CA3 pyramidal neurons in control and AD cultures after DTX-K application (grey: control cultures, orange: AD cultures). Note that the two curves are superimposed. Middle, Rheobase of CA3 pyramidal neurons in control and AD cultures after DTX-K application (grey: control cultures, orange: AD cultures). Note that the rheobases are not statistically different. Right, Input/output curves gain in control and AD cultures after DTX-K application (grey: control cultures, orange: AD cultures). Note that the gains are not statistically different.

**Supplementary Figure 3:**
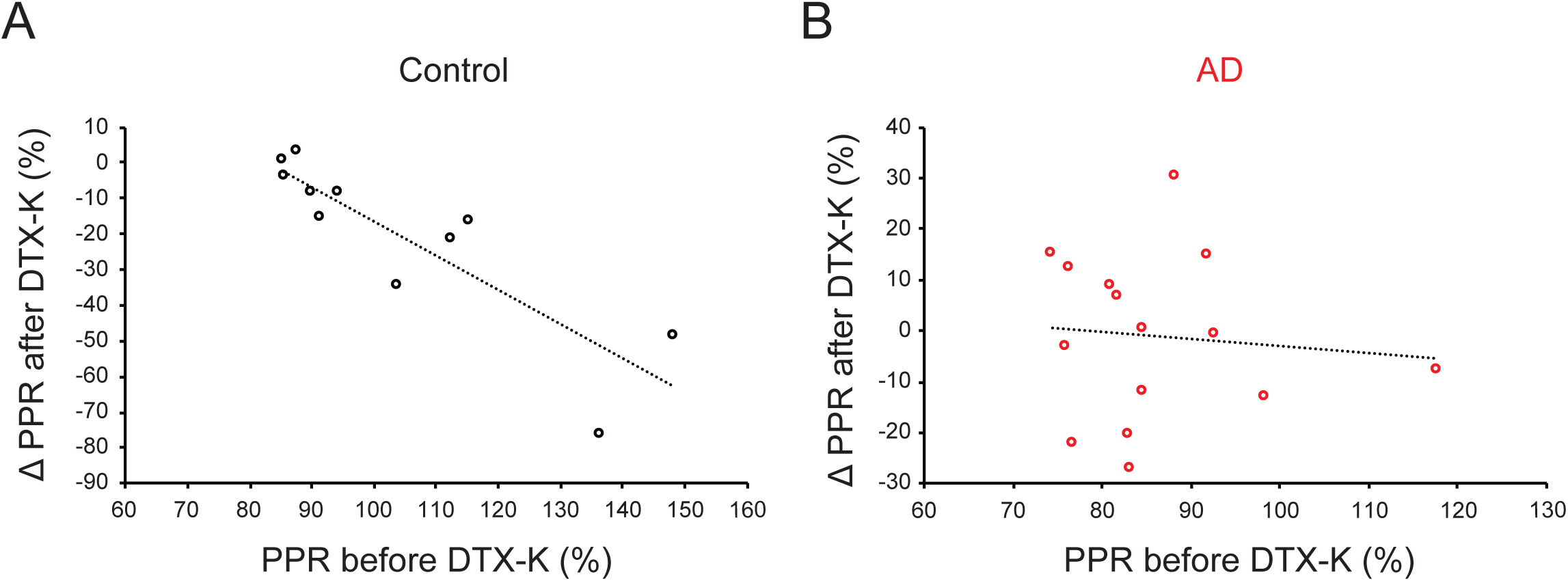
Synaptic connections with small release probability are more sensitive to K_v_1.1 blockade in control but not in AD cultures. Left, correlation between PPR before DTX-K and DTX-K effect on the PPR in control cultures (y = -0.9616x + 70.855; R^2^ = 0.7496). Right, absence of correlation between PPR before DTX-K and DTX-K effect on the PPR in AD cultures (y = -0.1347x + 10.421; R^2^ = 0.0086).

**Supplementary Figure 4:**
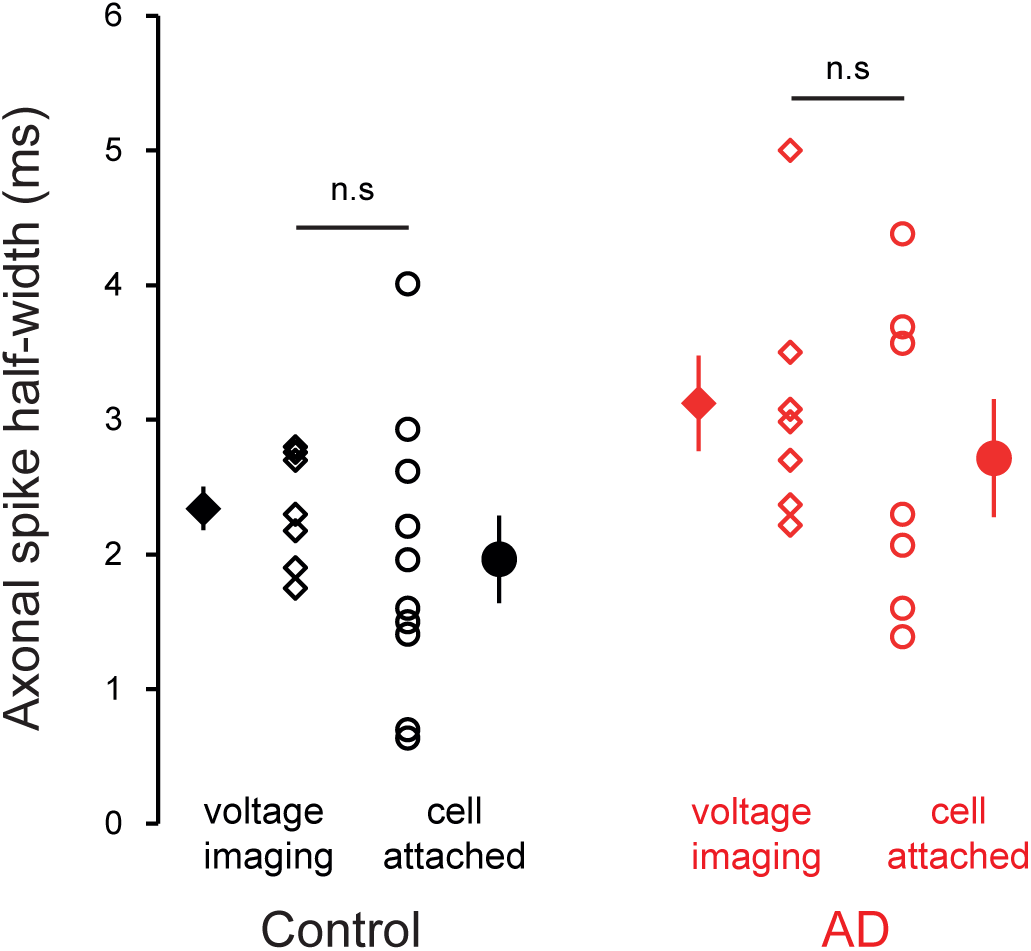
Similar axonal spike half-width found by voltage-imaging or cell-attached recordings. The values of axonal spike half-width measured by voltage-imaging or cell-attached recordings are not statistically different in control cultures (black). The values of axonal spike half-width measured by voltage-imaging or cell-attached recordings are not statistically different in AD cultures (red).

